# Repetitive subconcussive head impacts and changes in sensory processing for balance control

**DOI:** 10.1101/648857

**Authors:** Jaclyn B. Caccese, Fernando V. Santos, Eric Anson, Thomas A. Buckley, Felipe Yamaguchi, Mariana Gongora, John J. Jeka

**Affiliations:** University of Delaware, Department of Kinesiology and Applied Physiology, 540 S. College Ave., Newark, DE 19713, USA; University of Rochester, Department of Otolaryngology, 2365 S. Clinton Ave., Suite 200, Rochester, NY 14618, USA

**Keywords:** concussion, mild traumatic brain injury (mTBI), postural control, sensorimotor, vestibular

## Abstract

**Background:** Repetitive subconcussive head impacts (RHI) may be associated with current and future detrimental neurological effects. However, the effects of RHI on sensory processing for balance control is unknown and may have significant clinical implications if athletes are still participating in sport despite impairments.

**Research Question:** Are there changes in sensory processing for balance control during standing and walking following RHI?

**Methods:** Thirty healthy, adult, amateur soccer players (15 males, 15 females, 21.8±2.8 years, 69.9±11.5 kg, 171.4±8.2 cm) volunteered for the standing balance assessment (i.e., experiment 1). A distinct cohort of twenty healthy, adult, amateur soccer players (10 males, 10 females, 22.3±4.5 years, 70.0±10.5 kg, 170.5±9.8 cm) volunteered for the walking balance assessment (i.e., experiment 2). We used a repeated measures design across three time points (pre-heading, 0-hours post-heading, 24-hours post-heading). Participants were randomly assigned to an experimental (i.e., performed 10 soccer headers in 10 minutes) or a control group (i.e., did not perform any soccer heading between sessions). In experiment 1, participants stood in a virtual reality cave while experiencing simultaneous perturbations to their visual, vestibular, and proprioceptive systems. In experiment 2, participants walked blindfolded along a foam walkway and experienced a vestibular perturbation on the second heel strike of the right foot. Changes in sensory processing were assessed using repeated measures ANOVAs.

**Results:** There were no changes in sensory reweighting during experiment 1 and no changes in balance responses to the vestibular stimulus in experiment 2.

**Significance:** Although the cumulative effects of RHI may result in later-life cognitive, behavioral, and mood impairments, a single episode of repeated soccer headers does not appear to be associated with acute impairments in sensory processing for balance control.

## Introduction

Chronic Traumatic Encephalopathy (CTE) is a neurodegenerative disease thought to be caused by repetitive brain trauma that can occur during contact sports and military participation.[1] While initially reported in individuals with history of multiple concussions, or mild traumatic brain injuries, CTE has more recently been identified in football players with no history of diagnosed concussions, but with history of exposure to repetitive subconcussive head impacts (RHI).[1] A subconcussive head impact is a mild head impact that does not result in acute clinical signs or symptoms of neurological dysfunction, but may cause neurological deficits when sustained repeatedly.[2] Biomechanical investigations of head impacts in American football and soccer have indicated that athletes at the collegiate level sustain high numbers of RHI ranging from several hundred to well over 1000 during the course of a season.[3–6] Yet, the effects of RHI on neurological function are poorly understood.[2,7–10]

Quantifying changes in neurological function following RHI has several challenges. Clinical measures may not be sensitive enough to identify changes in neurological function following RHI. Though neuroimaging (e.g., functional MRI, DTI) can identify functional impairments and changes in white matter microstructure, we do not understand the clinical implications of these measures.[11–15] Moreover, on-field RHI exposure varies greatly in terms of magnitude, timing, type, and location of impact.[3–6] Variability in RHI exposure likely contributes to conflicting results across studies. To gain an understanding of the neurological deficits of RHI, we need sensitive measures of neurological function, while controlling for RHI exposure. An experimental paradigm using a series of soccer headers allows for control of RHI exposure.[16–21] This soccer heading paradigm has been used to understand the biomechanics of RHI and the effects of RHI on standing balance and biomarkers of head injury.[16–21] Humans require precise integration and modulation of visual, vestibular, and proprioceptive feedback to control balance during standing and walking. Visual and vestibular systems are vulnerable to disruption from concussion and may also be vulnerable to disruption from RHI.[22] Probing these sensory systems while standing and walking may allow us to identify subtle changes in neurological function following RHI. Hwang and colleagues developed an experimental paradigm to understand how sensory feedback is dynamically reweighted so that overall feedback remains suited to stabilizing upright stance.[23] To examine sensory reweighting, they simultaneously perturbed visual (using virtual reality), vestibular (using galvanic vestibular stimulation, GVS) and proprioceptive (using vibration) sensory modalities. Both intra-modal and inter-modal sensory reweighting occurred as the amplitude of visual input increased. This experimental paradigm can be used to examine the association between RHI and sensory reweighting.

Sensory modulation to balance has also been demonstrated during walking. GVS can be applied at different phases during locomotion and with varying amplitudes with little discomfort to the subject and with reduced concerns of adaptation.[24–26] Healthy adults respond to GVS applied during walking using a coordinated series of balance mechanisms.[27,28] Assuming that the perturbation occurs at heel strike, healthy adults sequentially use three mechanisms: lateral ankle roll followed by foot placement change and finally modulation of push off in the trailing leg.[27,28] The lateral ankle mechanism shifts the stance foot center of pressure (CoP) in the direction of the perceived fall by ~2.5 mm.[27,28] Foot placement modulation shifts the swing foot of the following step by ~15 mm in the same direction.[27,28] Finally, push off modulation adjusts the plantarflexion angle of the trailing leg to move the center of mass (CoM) in the direction opposite of the perceived fall.[28] Using these methods, this study aims to examine whether RHI affects sensory processing and subsequent balance control during standing and walking. We hypothesized immediately following the soccer heading paradigm individuals would have 1) impairments in sensory reweighting during standing and 2) diminished balance responses to GVS during walking.

## Methods

### Participants

Thirty soccer players (14 females, 21.8±2.8years, 69.9±11.5kg, 171.4±8.2cm) volunteered for the standing balance assessment (i.e., Experiment 1). A distinct cohort of twenty soccer players (10 females, 22.3±4.5years, 70.0±10.5kg, 170.5±9.8cm) volunteered for the walking balance assessment (i.e., Experiment 2). Potential participants were 18-35 years old with at least 5 years of soccer heading experience. Exclusion criteria included: head, neck, or lower extremity injury in the previous six months; history of balance problems; taking medications affecting balance; neurological disorders; unstable cardiac or pulmonary disease; goalkeepers due to lack of routine soccer heading practice. The university institutional review board approved the study and participants provided written informed consent. Participants in each experiment were randomly assigned to a heading group (EXP) or control group (CON). The EXP performed a soccer heading paradigm (described below), while the CON did not perform soccer heading.

### Experimental Design

We used a repeated measures design across three time points.[21] At each time point, participants completed either a standing balance assessment or a walking balance assessment (described below). The pre-heading session (PRE) was a baseline measurement. After 24h, the EXP performed 10 headers. The balance assessments were repeated immediately following the heading (POST-0h) and 24h later (POST-24h).

### Soccer Heading Paradigm

We used a soccer heading paradigm as an *in-vivo* model of repeated mild head impact.[17] We projected soccer balls (size 5, 450g, 8psi) using a JUGS soccer machine (JUGS Sports, Tualatin, OR); the initial velocity was 11.2m/s, the angle of projection was 40°, and the distance to the participant was 12m.[16–21] The EXP performed 10 headers in 10 minutes, while CON did not perform soccer heading.

### Experiment 1 - Standing Balance Assessment

We used an established paradigm for assessing sensory reweighting for standing balance (Figure 1).[23] Participants stood in a virtual reality cave (Bertec Corporation, Columbus, OH, USA), while experiencing simultaneous perturbations to visual, vestibular, and proprioceptive systems. Participants experienced four conditions: low amplitude visual scene translation with vibration and GVS (LVG), low amplitude visual scene translation without vibration but with GVS (LG), high amplitude visual scene translation with vibration and GVS (HVG), and high amplitude visual scene translation without vibration but with GVS (HG). Five trials for each condition were presented in random order for a twenty trials total. Trials lasted 135s.

**Figure 1.**
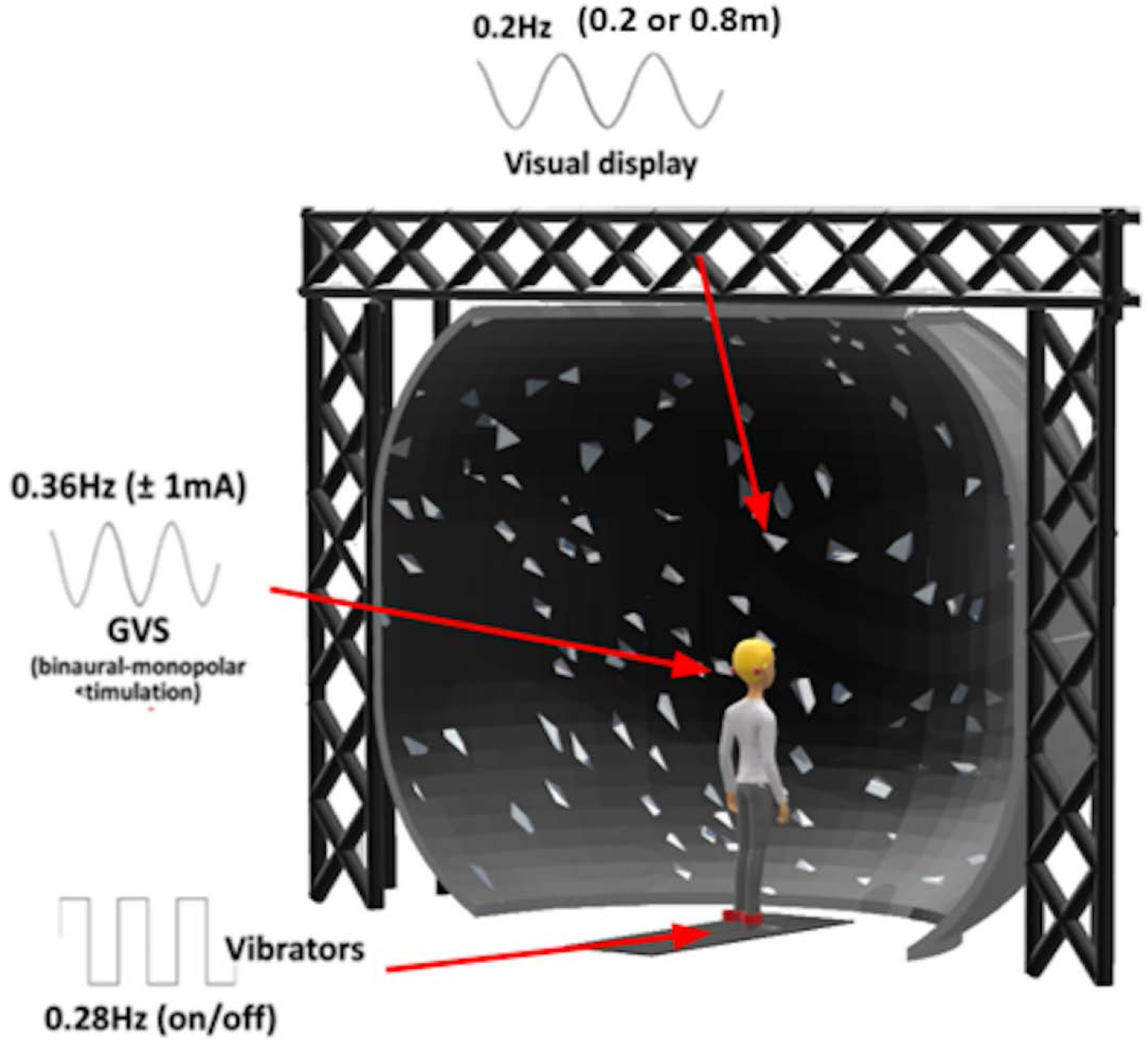
Set up for the standing balance assessment. Participants stood in a virtual reality while experiencing simultaneous perturbations to their visual, vestibular, and proprioceptive systems.

**Figure 2.**
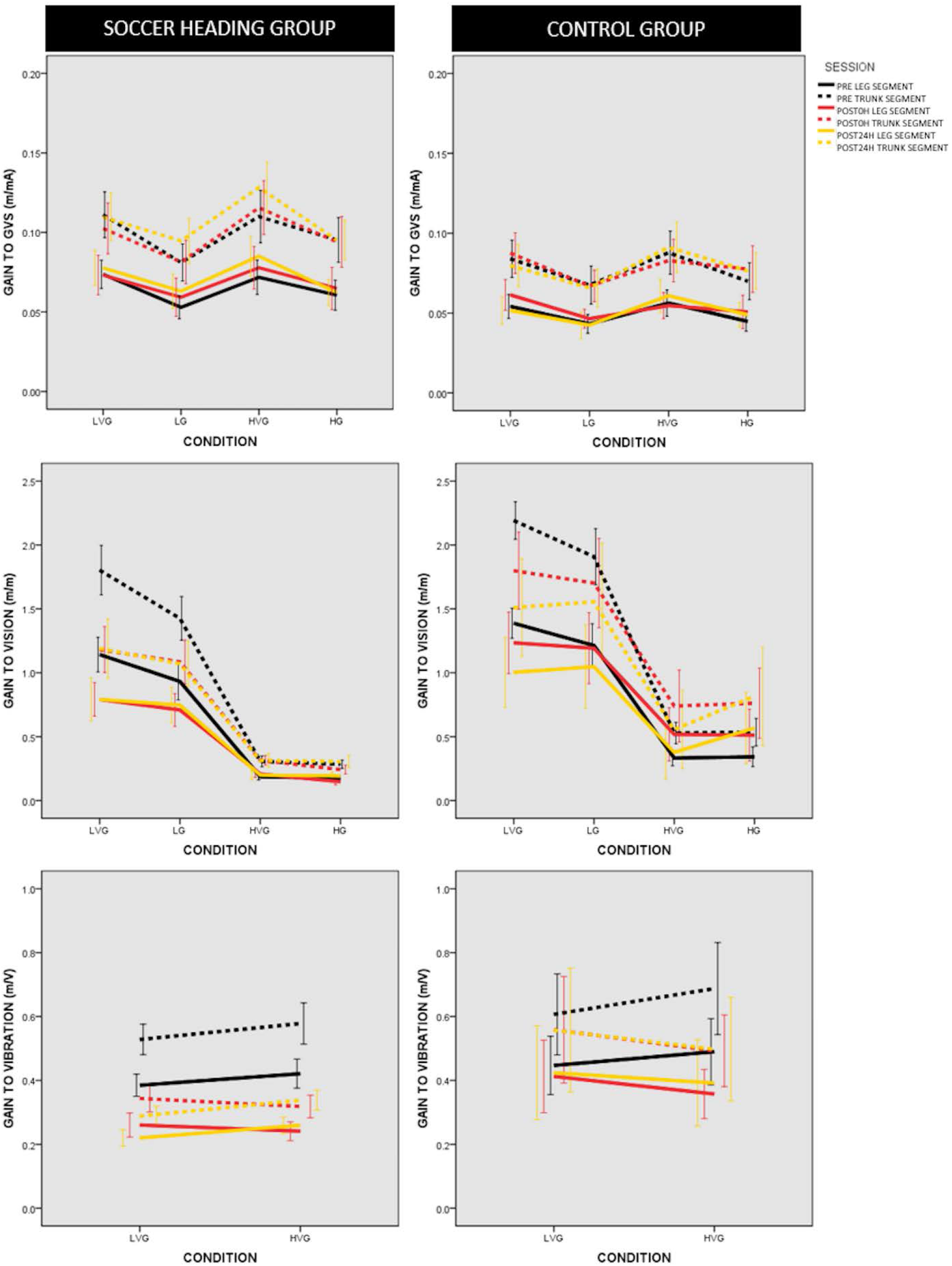
AP Leg (solid lines) and trunk (dashed lines) displacement gain to vision, gain to GVS, gain to vibration during the standing balance assessment. Lines represent group means ± standard error. Participants experienced four conditions: low amplitude visual scene translation with vibration and GVS (LVG), low amplitude visual scene translation without vibration but with GVS (LG), high amplitude visual scene translation with vibration and GVS (HVG), and high amplitude visual scene translation without vibration but with GVS (HG).

The visual scene consisted of 500 randomly distributed white pyramids on a black background, which translated sinusoidally at 0.2Hz in the anterior-posterior (AP) direction at two different amplitudes (0.2m and 0.8m for the low and high vision conditions, respectively). The proprioceptive sensory perturbation consisted of bilateral Achilles’ tendon vibrations from two custom vibrators in a 1mm displacement square-wave periodic stimulus at 0.28Hz. The vestibular sensory perturbation consisted of binaural-monopolar ±1mA sinusoidal GVS signal at 0.36Hz using a neuroConn DC-Stimulator Plus (neuroCare Group, Munchen, Germany). Uncorrelated frequencies were chosen for each stimulus so independent responses to each sensory modality could be determined.

Bilateral kinematics were collected at 120Hz using a twelve-camera motion analysis system (Qualisys, Goteborg, Sweden) and a marker set consisting of the temple, acromion, greater trochanter, lateral femoral condyle, lateral malleolus and first metatarsal head. The leg segment was defined by the AP displacement of the hip and ankle markers and the trunk segment by the AP displacement of the shoulder and hip markers. Gains and phases to each sensory modality were calculated. Gain is the amplitude of the output divided by the amplitude of the input at each driving frequency, calculated by the absolute value of the frequency response function, which is the cross spectral density divided by the power spectral density of the input.[23] Phase is a measure of the temporal relationship between the input and output; the output may lead the input or lag behind it.[23]

### Experiment 2 - Walking Balance Assessment

Participants walked blindfolded along a 2-inch closed-cell foam walkway centered over two force plates (AMTI, Watertown, MA, USA). Participants initiated gait with their right foot and took six steps until they were instructed to stop. Binaural, bipolar GVS was delivered from two electrodes with 3.2cm diameter (Axelgaard Manufacturing Co., Ltd, Fallbrook, CA, USA), placed on the mastoid processes. GVS was triggered on the second heel strike of the right foot and continued for 600ms. When triggered, a custom-made LabVIEW program (National instruments Inc., Austin, TX, USA) generated an analog voltage, which the neuroConn DC-Stimulator Plus transformed into a square wave of 1 mA current. In the GVS condition, the anode (LEFT) and cathode (RIGHT) created a perceived fall to the RIGHT. This perceived fall to the RIGHT creates an actual fall to the LEFT as a result of the actively generated motor response designed to prevent the perceived fall.[27] In the control condition, NO stimulation was delivered. Participants randomly repeated each of the two conditions 40 times, a total of 80 trials. Trials were excluded if the participant did not have a complete step on the force plate.

Bilateral kinematics were recorded at 120Hz using a 6-degree of freedom marker set. Kinetic data were recorded at 1200Hz. Muscle activity was recorded at 1200Hz using wireless electromyography (EMG; Trigno Wireless System, Delsys Inc., Natick, MA, USA). All data were analyzed in Visual 3D (C-Motion, Inc., Germantown, MD, USA). Kinematic and kinetic data were filtered at 6 and 25Hz, respectively, using a zero-lag, low-pass Butterworth filter, and were analyzed from right heel strike (RON) to right toe off (ROFF). The EMG data were rectified and smoothed using a 50ms moving RMS. Variables for the three balance mechanisms were computed, including the stepping mechanism, the lateral ankle mechanism, and the push off mechanism.[28] Each of three balance mechanisms is described by a functional outcome measure, corresponding joint kinematics, and representative muscle activity (Table 1).[28] For all measures, the mean of the control trials was subtracted from each of the stimulus trials to estimate the response to the stimulus.[27,28] The average maximum activation across control strides was used to normalize EMG.

**Table 1.**
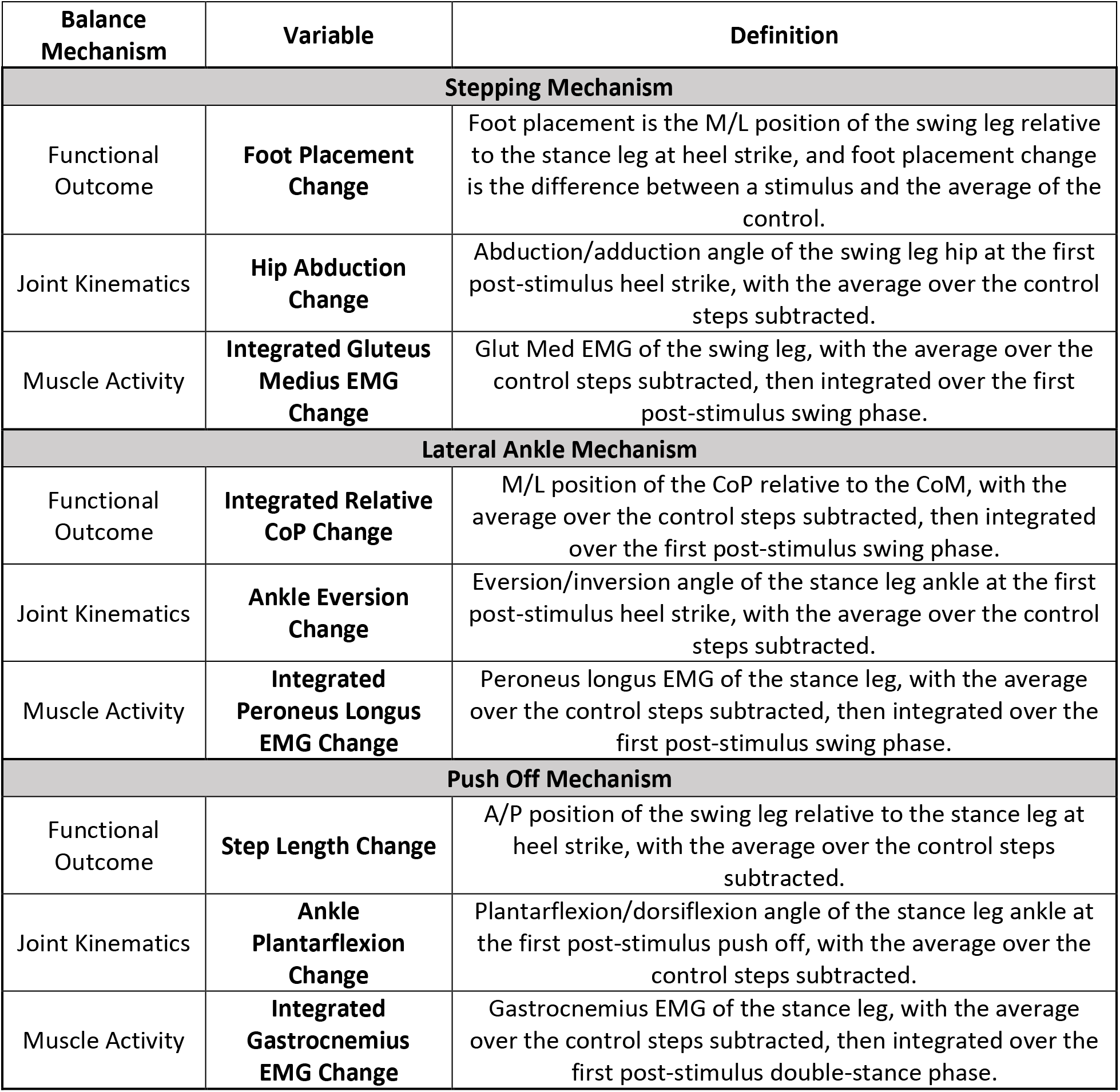
Definition of mechanisms used for balance control during walking.^30^

### Statistical Analysis

Repeated measures ANOVAs were used to compare mean response variables between groups (i.e. EXP vs. CON) across different time points (i.e. PRE, POST-0h, POST-24h). Outcome measures included gains and phases to each modality for Experiment 1 and the balance response variables for Experiment 2. SPSS (IBM Corporation, Armonk, NY) was used for all statistical analyses; α=0.05.

## Results

### Standing Balance Assessment – Leg AP Displacement

There were no changes in leg gain to vision (i.e. sessionXgroup effect; F=0.798, *P*=0.455, η^2^=0.028), to GVS (F=0.246, *P*=0.782, η^2^=0.009), or to vibration (F=0.662, *P*=0.520, η^2^=0.023) (Figure 1). In addition, there were no changes in sensory reweighting (i.e. sessionXconditionXgroup effect; vision, F=0.430, *P*=0.858, η^2^=0.015; GVS, F=0.763, *P* =0.600, η^2^=0.027; vibration, F=0.430, *P*=0.653, η^2^=0.015) (Figure 1).

Leg segment displacements displayed phase leads of ~25 deg relative to GVS and of ~50 deg relative to vibration. Relative to the visual stimulus, leg phase was zero. There were no changes in leg phase to vision (i.e. sessionXgroup effect; F=0.961, *P*=0.389, η^2^=0.033), to GVS (F=0.427, *P*=0.654, η^2^=0.015), or to vibration (F=0.375, *P*=0.689, η^2^=0.013). In addition, there were no changes in phase across conditions (i.e. sessionXconditionXgroup effect; vision, F=0.614, *P*=0.719, η^2^=0.021; GVS, F=1.238, *P*=0.289, η^2^=0.042; vibration, F=0.153, *P*=0.858, η^2^=0.005).

### Standing Balance Assessment – Trunk AP Displacement

There were no changes in trunk gain to vision (i.e. sessionXgroup effect; F=0.490, *P*=0.615, η^2^=0.017), to GVS (F=0.205, *P*=0.815, η^2^=0.007), or to vibration (F=0.624, *P*=0.539, η^2^=0.022). In addition, there were no changes in sensory reweighting (i.e. sessionXconditionXgroup effect; vision, F=0.395, *P*=0.881, η^2^=0.014; GVS, F=0.906, *P*=0.492, η^2^=0.031; vibration, F=0.761, *P*=0.472, η^2^=0.026).

Trunk segment displacements displayed phase leads of ~30 deg relative to GVS ~45 deg relative to vibration. Relative to the visual stimulus, trunk phase was zero. There were no changes in trunk phase to vision (i.e. sessionXgroup effect; F=1.257, *P*=0.292, η^2^=0.043), to GVS (F=0.656, *P*=0.523, η^2^=0.023), or to vibration (F=0.910, *P*=0.408, η^2^=0.031). In addition, there were no changes in phase across conditions (i.e. sessionXconditionXgroup effect; vision, F=0.636, *P*=0.701, η^2^=0.022; GVS, F=0.319, *P*=0.926, η^2^=0.011; vibration, F=1.247, *P*=0.295, η^2^=0.043).

### Walking Balance Assessment

There were no significant groupXtime interactions for any of the balance response variables (lateral foot placement, F=0.563, *P*=0.574, η^2^=0.030; hip abduction, F=0.038, *P*=0.963, η^2^=0.002; integrated gluteus medius EMG, F= 0.537, *P*=0.589, η^2^=0.029; integrated relative CoP, F=0.311, *P*=0.734, η^2^=0.017; ankle eversion, F=1.094, *P*=0.346, η^2^=0.057; integrated peroneus longus EMG, F=0.305, *P*=0.739, η^2^=0.017; step length, F=0.909, *P*=0.412, η^2^=0.048; ankle plantar flexion, F=0.610, *P*=0.549, η^2^=0.033; integrated gastrocnemius EMG, F=0.547, *P*=0.583, η^2^=0.029) (Table 2).

**Table 2.** Balance response variables during the walking balance assessment for each group (i.e., CON, EXP) across time (i.e., PRE, POST-0h, POST-24h). Data represent mean ± standard deviation.

| OUTCOME | GROUP | PRE | POST-0h | POST-24h |
| --- | --- | --- | --- | --- |
| **Stepping Mechanism** |  |  |  |  |
| **Foot Placement Change (m)** | **CON** | 0.030±0.017 | 0.030±0.014 | 0.025±0.017 |
| | **EXP** | 0.034±0.024 | 0.033±0.018 | 0.022±0.009 |
| **Hip Abduction Change (deg)** | **CON** | -1.146±0.677 | -1.202±0.820 | -0.938±0.714 |
| | **EXP** | -0.753±0.813 | -0.869±1.127 | -0.502±0.801 |
| **Integrated Gluteus Medius EMG Change (normalized\*s)** | **CON** | -0.002±0.007 | -0.005±0.003 | -0.005±0.010 |
| | **EXP** | -0.002±0.009 | -0.001±0.006 | -0.004±0.005 |
| **Lateral Ankle Mechanism** |  |  |  |  |
| **Integrated Relative CoP Change (m\*s)** | **CON** | 0.002±0.002 | 0.001±0.002 | 0.002±0.001 |
| | **EXP** | 0.003±0.003 | 0.002±0.002 | 0.003±0.002 |
| **Ankle Eversion Change (deg)** | **CON** | 5.606±2.801 | 5.467±3.183 | 4.217±2.604 |
| | **EXP** | 8.199±2.608 | 6.339±1.710 | 5.510±3.030 |
| **Integrated Peroneus Longus EMG Change (normalized\*s)** | **CON** | -0.057±0.043 | -0.058±0.049 | -0.062±0.049 |
| | **EXP** | -0.089±0.052 | -0.080±0.047 | -0.076±0.046 |
| **Push Off Mechanism** |  |  |  |  |
| **Step Length Change (m)** | **CON** | 0.007±0.012 | 0.007±0.014 | 0.008±0.007 |
| | **EXP** | 0.011±0.007 | 0.011±0.012 | 0.006±0.008 |
| **Ankle Plantarflexion Change (deg)** | **CON** | 0.001±0.009 | 0.001±0.005 | 0.001±0.007 |
| | **EXP** | -0.002±0.013 | -0.005±0.009 | -0.002±0.008 |
| **Integrated Gastrocnemius EMG Change (normalized\*s)** | **CON** | -0.215±3.073 | -0.284±2.456 | -0.239±1.849 |
| | **EXP** | -1.353±2.955 | -1.648±2.455 | -0.532±3.673 |

## Discussion

While the acute effects of concussion have been characterized in the literature, the effects of RHI are poorly understood.[2,7–10] We examined the association between RHI and sensory processing for balance control during standing and walking. We hypothesized immediately following the soccer heading paradigm individuals would have 1) impairments in sensory reweighting during standing and 2) diminished balance responses to GVS during walking. Our findings did not support our hypotheses; there were no changes in sensory reweighting during standing or balance responses to GVS during walking. Despite concerns that RHI exposure through routine contact sport participation may result balance disturbances, our results do not support this association. Our findings are consistent with previous work suggesting repeated soccer heading has no effect on postural stability;[29–31] however, it should be noted that other studies were able to challenge the postural control system enough to elicit changes in standing balance.[16,21] We speculate contradictory results across studies may be attributed to the complexity/challenge of the balance task. For example, Haran and colleagues assessed participants in 6 virtual environment (VE) conditions: (1) stationary VE with stationary support surface, (2) in the dark with rotating support surface, (3) rotating VE with stationary support surface, (4) rotating VE with rotating support surface, (5) in the dark with stationary support surface, (6) stationary VE with rotating support surface, and *only* observed differences in condition 4, a particularly destabilizing condition.[16] Similarly, Hwang and colleagues noted decreased gain to GVS following RHI when participants were standing blindfolded on a foam surface; therefore, eliminating visual feedback and reducing proprioceptive feedback.[21] These studies elicited changes in standing balance, although translatability of these assessments to “real-world” balance tasks during which full sensory information is available remains an open question.

Despite growing public concern regarding the potentially adverse effects of RHI, our results taken in the context of previous literature suggest there are few immediate changes in behavioral measures following RHI, notwithstanding changes in neuroimaging and blood biomarkers.[10,32,33] In both animal and clinical studies, RHI have associated with Blood Brain Barrier dysfunction, abnormal neuro-metabolic and neuroinflammatory processes, as well as Tau aggregation.[10] In contrast, behavioral changes appear to be minimal. Caccese and colleagues used a multimodal clinical battery to examine changes in balance and neurocognitive function in collegiate women’s soccer and men’s American football players.[33] There were no changes in any of the measures from pre-season to post-season.[33] Additionally, Buckley and colleagues assessed changes in single-task and dual-task balance in American football players and non-contact athletes throughout a season; there were no changes for any of the dynamic balance tasks.[32] These findings are consistent with most studies which fail to identify short-term clinical differences related to RHI; however, it is possible changes in neuroimaging and blood biomarkers manifest into long-term clinical impairments.

### Experiment 1

In healthy adults, leg and trunk gain relative to vision decreases from low vision conditions to high vision conditions, which is intramodal reweighting.[23] In other words, as the visual scene amplitude increases, reliance on vision decreases (down-weighting). Similarly, leg and trunk gain relative to GVS stimulus and to vibration stimulus increases when visual amplitude increases, intermodal reweighting, to compensate for visual down-weighting. In our cohort, we see clear evidence of both intramodal and intermodal reweighting (Figure 1), suggesting that these individuals can perceive changes in sensory stimuli and generate appropriate sensorimotor responses. Although our standing balance assessment evaluated the dynamic modulation of three modalities, we did not observe changes in sensory reweighting after RHI. Our results differ from those of Hwang and colleagues, who reported diminished gain to GVS following soccer heading.[21] Importantly, the current protocol examined *sensory reweighting*, all sensory systems were perturbed simultaneously; whereas, Hwang and colleagues limited the use of vision with a blindfold and made proprioception constantly unreliable with a foam surface.[34,35] Perhaps when other sensory information is available (as in this experiment) participants can reweight sensory information to maintain upright stance, but under more extreme challenges when only vestibular information is available, participants experience diminished vestibular processing.

### Experiment 2

In response to GVS stimulation during walking, healthy adults sway laterally in the direction away from the perceived fall.[27,28] Several corrective mechanisms are used to regain balance following this type of perturbation, including the lateral ankle mechanism, stepping mechanism, and push off mechanism.[28] This study was the first to quantify corrective balance mechanisms during GVS perturbed walking following RHI exposure. Although we observed small, systematic indications of each walking balance mechanism (i.e., participants responded to GVS perturbation), we did not observe changes in balance response variables following RHI (Table 2). There was substantial variability in how individuals use balance mechanisms (Table 2), which is consistent with previous research of individuals walking on a foam surface.[36] Although these complementary mechanisms allow for dynamic control in response to balance perturbations, the variability within and across individuals likely masked changes.

### Limitations

Our interpretations are limited to effects of RHI on balance control during standing and walking and do not preclude changes in other neurological measures, such as cognition and mood. Future work should incorporate a multimodal battery of assessments. Although our soccer heading paradigm was designed to control RHI, 10 soccer headers may be insufficient to elicit balance deficits. RHI may cause neurological deficits that manifest over a career of repetitive head trauma and not after a single RHI exposure.

## Conclusion

We examined whether sensory reweighting during standing and balance responses to GVS during walking were altered after controlled RHI. Although the cumulative effects of RHI may result in later-life neurological deficits, a single episode of RHI was not associated with impairments in sensory processing for balance control. These findings have significant clinical implications; any impairments in balance as a result of RHI may predispose athletes to subsequent injury, so it is reassuring to find that there were no deficits in sensory processing for balance control during standing or walking.

